# Global statistical models of protein coevolution reveal higher-order sectors beyond those obtained from structure alone

**DOI:** 10.1101/2022.05.27.493723

**Authors:** Carina Shiau, Haobo Wang, Young Lee, Sergey Ovchinnikov

## Abstract

Recent methods have shown promise in using pairwise sequence coevolution predictions to illuminate physical interactions and functional relationships between pairs of protein residues. As a result, there has been an increased interest in identifying higher-order correlations between sequence positions in an effort to further understand how the multiple sequence alignment (MSA) encodes the conserved biological properties of a protein family. To this end, we propose a robust and generalizable spectral clustering model that can extract interconnected networks of coevolving residues – termed “protein sectors” – using pairwise sequence coevolution predicted by a global statistical model. We assess the statistical and evolutionary origins for protein sectors extracted from the MSA for 120 protein families. We show that protein sectors are extracted from a subset of densely connected components in the sequence coevolution matrix, many of which are not present in the pairwise residue contact graph that is constructed from the protein crystal structure, revealing the existence of networks of paired residues that are not necessarily in direct physical contact but are nonetheless evolutionarily coupled. We found that protein sectors form structurally connected entities in three-dimensional space, despite sector identification being independent of protein crystal structure. Interestingly, protein families with high structural similarity do not share similar protein sectors, suggesting that nuances in the sequence coevolution matrix can differentiate between the evolutionary histories of structurally-related protein families.

## 2 Introduction

The rapid progress in DNA sequencing techniques and the inclusion of metagenome sequence data has generated more than 3.6 *×*107 known protein sequences for 14000 protein families [1], with the number of sequences increasing by as much as 100-fold for some families [2]. In parallel with this growth of sequence databases, there has been a plethora of efforts to use sequence data alone to extract information about the biochemical and structural features of a protein that may be under evolutionary pressure [1, 3, 4, 5, 6].

One measure has been single-residue conservation. In the absence of selective pressures, the distribution of amino acids at a given position across the multiple sequence alignment is expected to be random and demonstrate low conservation [4]. However, amidst the functional and structural constraints that act on the evolution of a protein, certain positions have proven to be highly conserved and easily detectable in the multiple sequence alignment of a given protein family. Residue mutations at these positions would likely disrupt protein function, which could be due to the residue’s role in the enzymatic active site or in the structural integrity of the protein [1]. Thus, single-residue conservation measures the strength of evolutionary interference at a given position and therefore the biological significance of a given residue [2, 4].

However, evolutionary pressure does not only result in single-site conservation. Due to co-adaptation between pairs of coevolving residues, a mutation in one residue may alter the fitness landscape of another interacting residue. Compensatory mutations can preserve the integrity of a protein [1, 4]. In other words, the evolution of a protein is not independent but instead inter-dependent. Thus, we turn to analyzing molecular coevolution between residues. Early approaches to extract coevolution signals from the multiple sequence alignment include Mutual Information (MI) [7], Coevolution Analysis using Protein Sequences (CAPS) [8], and Statistical Coupling Analysis (SCA) [9]. However, these methods often do not distinguish between direct couplings and indirect correlations that arise from chains of direct couplings [10, 11]. Later methodological improvements seek to achieve separation between direct and indirect couplings by using a global statistical model. Direct Coupling Analysis (DCA) [12, 13, 14] and Protein Sparse Inverse COVariance (PSICOV) [15] rely on estimating the inverse covariance matrix, while recent work with the GREMLIN Learning Algorithm show that a pseudo-likelihood-based approach can produce even more accurate results for a range of alignment sizes and protein lengths [10]. Indeed, these analyses of molecular coevolution have revealed crucial residue-residue interactions, often illuminating physical interactions or functional relationships between pairs of residues.

These findings motivate further analysis of higher-order correlations between sequence positions in an effort to understand how the multiple sequence alignment encodes the conserved biological properties of a protein family. Previous studies on one representative protein family (the S1A family of serine proteases) found nonrandom higher-order correlations in the SCA matrix between sequence positions that decompose the protein length into groups of coevolving amino acids, termed “protein sectors.” The identified protein sectors in the S1A family of serine proteases were physically connected in the tertiary structure. The protein sectors were also nearly statistically independent, meaning that the pairwise correlation entropies were approximately equal to the sum of individual correlation entropies. Further experimentation also led to the hypothesis that distinct protein sectors controlled distinct phenotypes of the protein, such as thermodynamic stability and catalytic activity [3, 4].

However, the validity of protein sectors has been heavily debated. Previous methods for sector extraction relied on SCA, which is unable to differentiate between indirect and direct couplings. This questions whether protein sectors are truly interconnected networks of direct couplings or whether protein sectors are simply the result of chains of indirect couplings. It is uncertain to what extent the composition of a protein sector is dependent on evolution as opposed to which statistical analysis method, parameters, and thresholds are employed [4]. To this end, we propose a robust and generalizable spectral clustering algorithm for protein sector extraction. The model utilizes sequence coevolution data predicted by a global statistical model that promotes a sparse network. Applying this model to 120 protein families, each containing over 1000 aligned sequences, we demonstrate that protein sectors are densely connected networks of coevolved residue pairs that are in spatial proximity in the three-dimensional crystal structure. Interestingly, protein families with high structural similarity do not share similar protein sectors. While sequence coevolution has often been used as a tool for structure prediction, this finding suggests that the sequence coevolution matrix may contain nuances that can help us to differentiate between structurally-related protein families, perhaps revealing an evolutionary history of conserved functional properties.

## 3 Results

### 3.1 Spectral decomposition and clustering of sequence coevolution data yields residue networks, or protein sectors

The multiple sequence alignment (MSA) for a protein family contains a reserve of single-residue conservation and pairwise coevolution information. Single-residue conservation measures the strength of evolutionary interference at a single residue site (indicated by the green residue in Figure 1A left), while pairwise coevolution measures the strength of evolutionary inference at residue-residue junctions (indicated by the yellow and blue residues in Figure 1A left). The GREMLIN (Generative REgularized ModeLs of proteINs) Learning Algorithm – a global statistical model – uses the MSA to estimate the sequence coevolution matrix, and average product correction (APC) is subsequently applied to disentangle structural and functional signals from background noise and phylogeny. The resultant AP-corrected coevolution matrix (Figure 1A center) contains higher-order correlations between sequence positions, which we term “protein sectors” (Figure 1A right).

**Figure 1:**
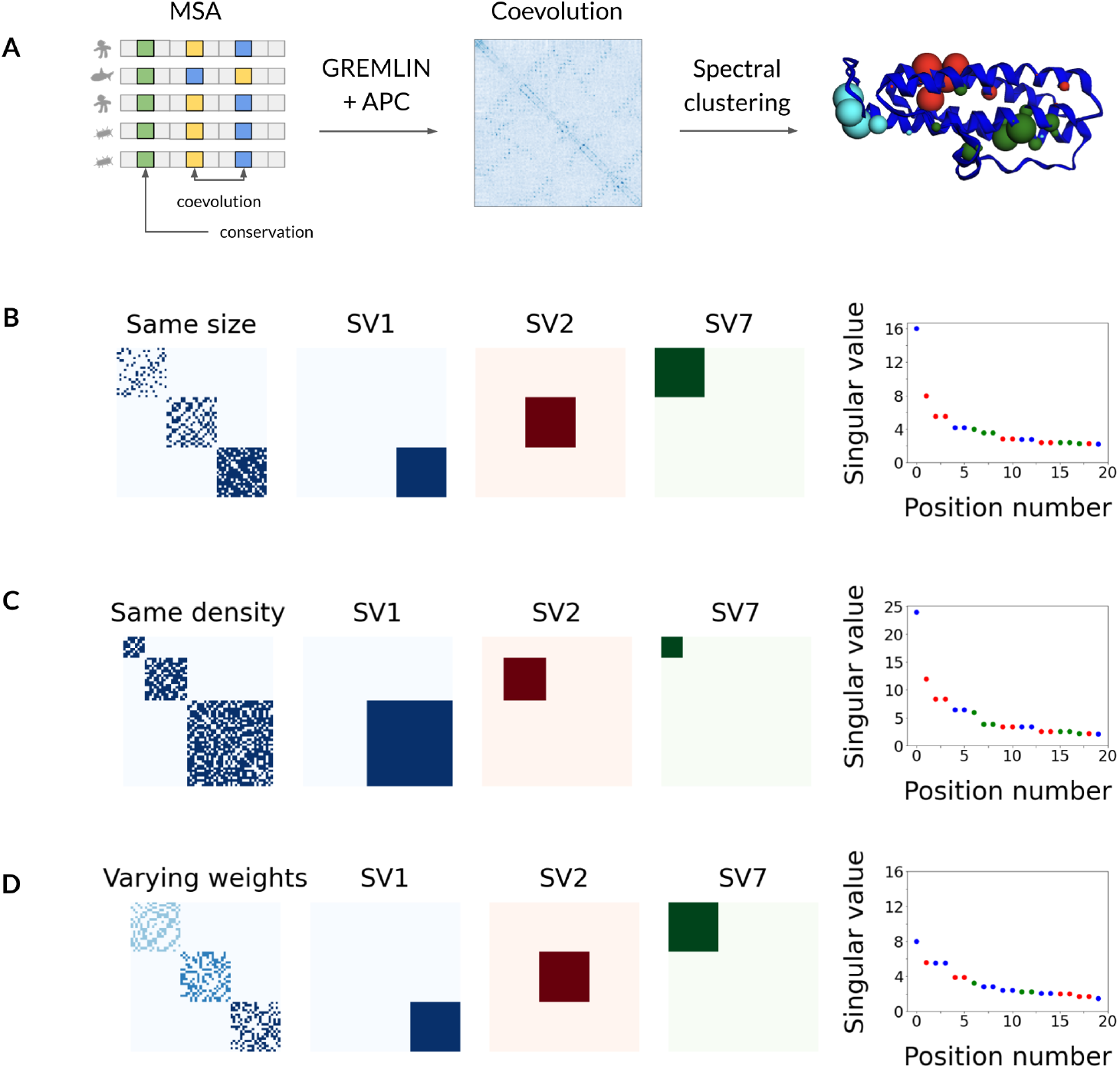
Singular value decomposition (SVD) can be leveraged to identify protein sectors by prioritizing extraction of more densely connected clusters and of larger residue networks. (A) Protein sector extraction model for the 3AK8 protein family employs spectral clustering and the multiple sequence alignment (MSA). The MSA (left) contains conserved (green squares) and coevolving (yellow and blue squares) residues. GREMLIN models the MSA to estimate the sequence coevolution matrix, which is adjusted via average product correction (APC). Spectral clustering on the AP-corrected coevolution matrix (center) yields 3 protein sectors, which are mapped onto the 3AK8 PDB structure as differently-colored spheres (right). Sphere size at a given position corresponds to the non-negative matrix factorization (NMF) probability that the residue belongs to a specific sector. (B-D) Spectral decomposition for toy examples with 3 clusters of decreasing sparsity (B), 3 clusters of increasing size and constant density (C), and 3 clusters of varying weights (D). For each example, we show the adjacency matrix (left); the product of first, second, the seventh largest singular vectors (SV) with their transpose (center); and the top 20 singular values (right). The color for each singular value (right) denotes which cluster (left) its corresponding singular vector recreates (center).

Protein sectors are densely connected components of the sequence coevolution matrix that indicate interconnected networks of coevolving residues. We use singular value decomposition (SVD) to develop a spectral clustering model that can predict the protein sectors from the multiple sequence alignment of a given protein family. The full model is described in “Methods and Materials” and, in detail, in “Supplementary Methods”.

We use a toy model to demonstrate how SVD is capable of extracting protein sectors from the sequence coevolution matrix. Figures 1B - 1D present 3 toy examples in which we simulate adjacency matrices that contain clear residue networks. We present examples of networks that range in size, density, and weights. We want to extract only the networks that involve a larger number of residues, that are more densely connected, and that have heavier edge weights.

In Figure 1B left, we present an example with 3 clusters of decreasing sparsity, to which we decompose into singular values and singular vectors. The most densely connected cluster is described by the largest singular value and its corresponding singular vector. The largest singular value is 16 (Figure 1B right), which indicates the number of connections each residue has in the most densely connected cluster. The product of the largest singular vector with its transpose is a matrix that identifies the location of the most densely connected sector (Figure 1B center, blue). Similarly, the second most densely connected sector is described by the second largest singular value and its corresponding singular vector. The second largest singular value is 8 (Figure 1B right), which also corresponds to the number of connections in the second most densely connected sector. The product of the second largest singular vector with its transpose is a matrix that identifies the location of the second most densely connected sector (Figure 1B center, red). However, the most sparsely connected sector is not described until the seventh largest singular value (Figure 1B right) and its corresponding singular vector (Figure 1B center, green).

In Figure 1C left, we decompose another example with 3 clusters of increasing size but constant density. The smallest cluster is not described until the seventh singular value (Figure 1C right) and its corresponding singular vector (Figure 1C center, green), while the largest and second largest clusters were described in the first and second singular values, respectively. In Figure 1D, the cluster with the lightest edge weights is also not described until the seventh singular value, while the clusters with heavier edge weights are described in the first and second singular values. Through these 3 examples (Figure 1B - 1D), we illustrate that SVD can prioritize extracting more densely connected and larger clusters with heavier weights, enabling us to identify protein sectors from the sequence coevolution matrix.

### 3.2 Protein sectors form spatially connected entities in the protein crystal structure

The identification of protein sectors is entirely dependent on the statistical analysis of protein MSA without consideration of the three-dimensional protein crystal structure. Nonetheless, the protein sectors form compact structural entities, as shown in the protein structure from Figure 1A right.

We quantify the structural connectivity of protein sectors in Figure 2 by measuring the distance of a residue from its sector centroid with respect to its probability of being part of a given sector across 120 examples of protein families. We introduce the NMF probability (detailed in “Methods and Materials” and in “Supplementary Methods”) as a measure that is output from the sector extraction model to describe sector membership for a given residue. In Figure 2A, we observe that the distances from sector centroid decreases with respect to increasing NMF probabilities. In other words, a given residue *i* with a large NMF probability for sector *k* is more likely to be closer in three-dimensional distance to sector *k*’s centroid than another residue *j* with a smaller NMF probability for the same sector *k*. We also note that the mean distances (2A) and distributions of distances (2B) at each NMF probability range differ significantly from randomly-generated null sectors (Kolmogorov-Smirnov test).

**Figure 2:**
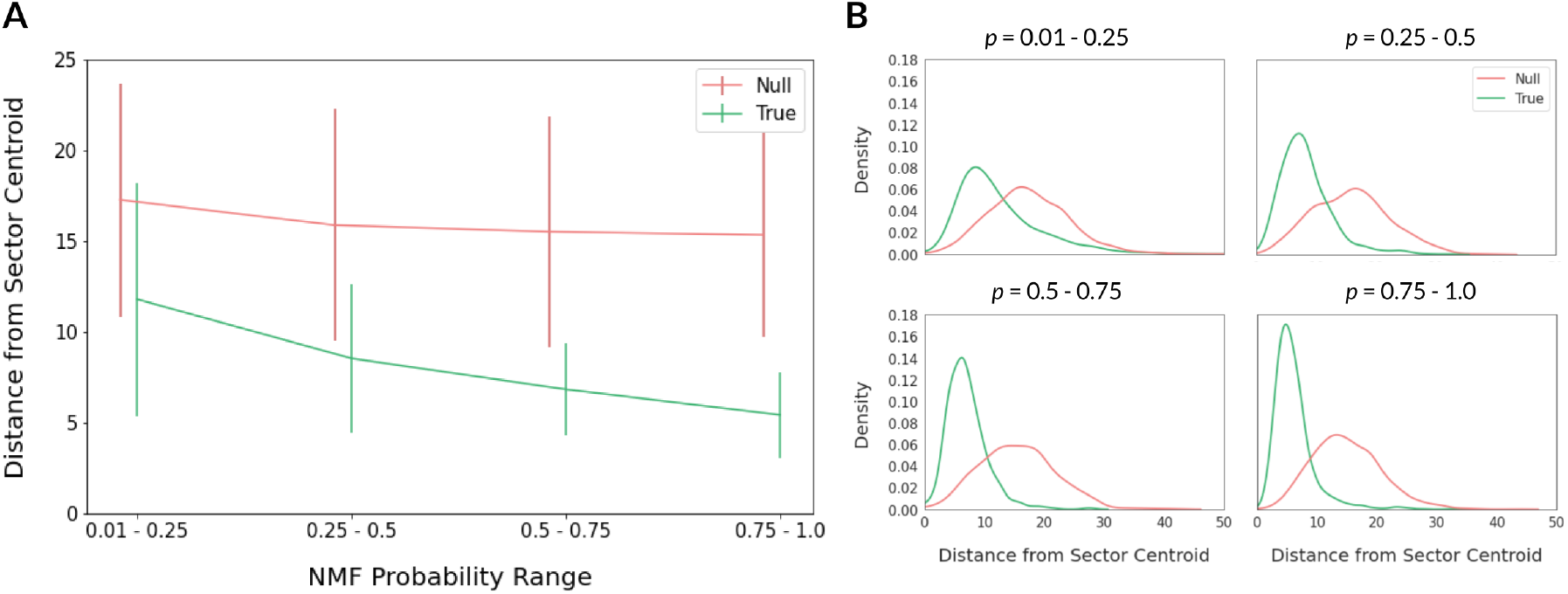
Protein sectors are physically connected in the three-dimensional protein structure. For each residue with non-negative matrix factorization (NMF) probability *≥* 0.01 of belonging to a certain sector, we calculate the distance of the residue in Angstroms from the sector’s centroid. (A) We compare residue distances from sector centroid at 4 ranges of NMF probabilities for true (green) versus null sectors (pink) across 120 proteins. Error bars show the standard deviation of the distances. (B) Distributions of residue distances from sector centroid at 4 ranges of NMF probabilities for true (green) versus null sectors (pink). True sectors are extracted using the spectral clustering model, while null sectors are randomly-selected positions from the protein sequence length.

Thus, the protein sectors are significantly more compact in physical space than what is expected for a set of randomly-selected residues.

### 3.3 Protein sectors form densely connected networks of coevolving residues, many of which are not in contact in the protein crystal structure

The sequence coevolution matrix has been found to accurately predict residue-residue contacts within a protein family. This correspondence, along with the structural connectivity of protein sectors (Section 3.2), lends itself to the following question: Are residue-residue contacts alone sufficient for sector extraction? We explore this question through 2 approaches that target the question from opposite ends: 1) determine whether extracted sectors form densely connected components in the contact map, and 2) perform spectral clustering on the contact map to extract “sectors”.

In the first approach, we thoroughly investigate the sectors of 4 protein families, shown in Figures 3A-D. For example, the spectral clustering model predicts 3 protein sectors for the 3AK8 protein family. These protein sectors can be interpreted as 3 graph networks, in which the graph nodes are residues with NMF probability*≥* 0.01 of sector membership and the graph edges are the connected components in the sequence coevolution matrix (Figure 3A, bottom). We can also create another graph using the same graph nodes but instead with graph edges that denote connected components in the protein contact map (Figure 3A, top). The protein contact map is an *L× L* similarity matrix in which all pairwise contacts with ConFind scores *cf≥* 0.01 are determined to be in contact in the three-dimensional crystal structure and indicated as a 1; otherwise, 0. When comparing the paired graph networks, notice how 2 of 3 graph networks with coevolution-based edges (left and right) form singular connected components (darker weight colors) across nearly all its nodes, while none of the 3 graph networks with contacts-based edges form singular connected components across the majority of nodes. Unlike the dense network formed by the coevolution-based edges (Figure 3A, bottom), the contacts-based edges only span a fraction of the nodes in a loose network (Figure 3A, top).

**Figure 3:**
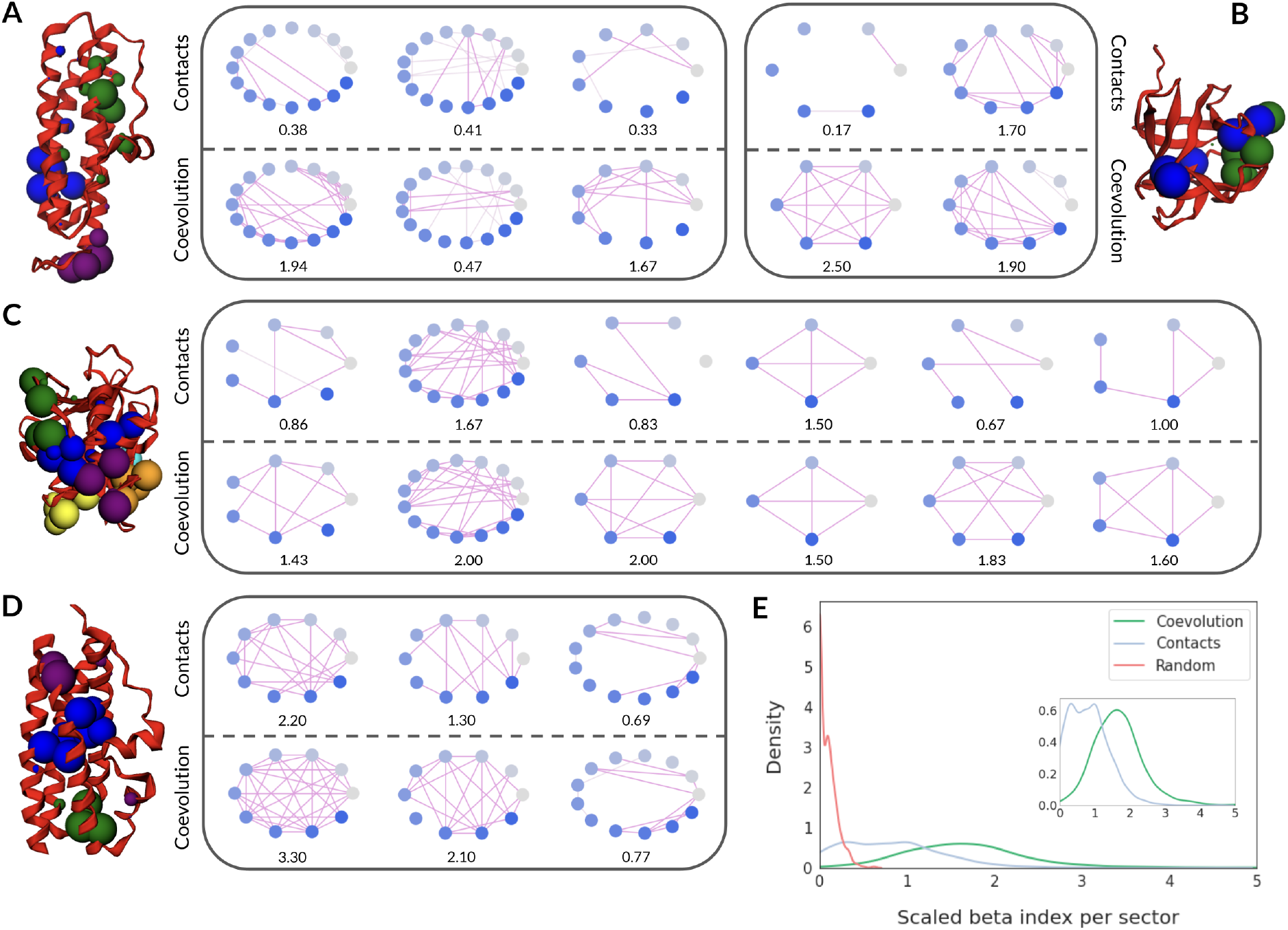
Protein sectors are densely connected components in the sequence coevolution matrix that could not have been extracted from the contact matrix of the protein crystal structure. (A-D) Graph networks and three-dimensional structures for the 3AK8 (A), 3P2H (B), 2BUE (C), and 3BT5 (D) protein families. For example, the spectral clustering model predicts 3 protein sectors for 3AK8, which are visualized in the upper 3 graph networks in (A) and again in the lower 3 graph networks in (A). Graph nodes are residues with NMF probability *≥* 0.01 for a particular protein sector. Edges in the lower 3 graph networks denote connected components in the sequence coevolution matrix, while the edges in the upper 3 graph networks denote connected components in the crystal structure-based contact matrix (residue pairs with ConFind scores *cf≥* 0.01 were determined as in-contact). Darker colored edges denote the largest connected component in the graph networks. The scaled beta index *β′* is noted under each graph network, indicating the product of its level of connectivity (beta index *β*) and the fraction of nodes that is participating in the largest connected component. (E) Comparison between coevolution-based scaled beta indices for contacts-generated sectors, coevolution-generated sectors, and randomly-generated sectors across 116 protein families. Here, the scaled beta index *β′* measures the connectivity of graph networks using the differentially-generated sectors as nodes and top signals from the sequence coevolution matrix as edges. The inset narrows in on the distributions for coevolution- and contacts-generated sectors for easier scaled beta index comparison.

We can quantify this difference by computing the scaled beta index (noted under each graph network in Figure 3A) for each graph. The scaled beta index for a graph is a product of the fraction of nodes participating in the largest connected network and the level of connectivity (beta index) of that network. Simple networks have scaled beta indices less than one, connected networks encompassing all nodes with one cycle have scaled beta indices equal to one, while complex networks encompassing all nodes have scaled beta indices greater than one. For the 3AK8 protein family, 2 of 3 graph networks (left and right) with coevolution-based edges (Figure 3A, bottom) have scaled beta indices much greater than 1. Given that most nodes are participating in the connected networks, this indicates that the sectors are densely connected, and almost every node is reachable from any other node. In contrast, all 3 graph networks with contacts-based edges (Figure 3A, top) have scaled beta indices much less than 1, indicating simple networks.

We can perform a similar analysis across 3 more protein families (Figures 3B-D). We observe that all graphs with coevolution-based edges form a densely connected network across all nodes with scaled beta indices greater than 1, with the exception of the third graph network in the 3BT5 protein family (Figure 3D). On the other hand, the majority of graph networks with contacts-based edges do not form singular connected components, and the majority do not have scaled beta indices greater than 1. Moreover, all graph networks with coevolution-based edges have stronger connectivity, i.e. greater scaled beta indices, than its corresponding graph networks with contacts-based edges, without exception.

Thus, we can conclude from the first approach to answering this section’s leading question that extracted protein sectors do not reliably form densely connected components in the protein contact map. We are unable to use the protein contact map to reconstruct the same sectors as those extracted from the sequence coevolution matrix. We can further support this conclusion through the second approach outlined above.

In the second approach, we would like to investigate whether “sectors” extracted from the contact map can recreate the protein sectors extracted from the sequence coevolution matrix. To this end, we compare the scaled beta indices of coevolution-generated protein sectors and contacts-generated protein sectors across 120 protein families (Figure 3E). Contacts-generated protein sectors are extracted by performing spectral clustering on the binary contact map (as determined by ConFind). We translate coevolution- and contacts-generated sectors into graph networks, for which we can compute the scaled beta indices. For equal metric comparison, all graph edges are composed of the top signals in the sequence coevolution matrix.

We observe distinct distributions between the scaled beta indices for ceovolution- and contacts-generated sectors (Figures 3E). The distribution for coevolution-generated sectors is centered between 1 and 2, while the distribution for contacts-generated sectors is centered below 1. We also included a negative control distribution for randomly-generated sectors, which was concentrated close to 0, as expected. These distributions further support our previous finding from the first approach that coevolution-generated sectors are densely connected components in the sequence coevolution matrix. The distributions also suggest that performing spectral clustering on the protein contact map cannot yield sectors that are densely connected in the sequence coevolution matrix.

The lack of overlap between coevolution- and contacts-generated sectors indicates that the spectral clustering model is not solely extracting coevolution signals that correspond to physical contacts in the three-dimensional protein crystal structure. The model is also reliant on signals in the sequence coevolution matrix that do not predict pairwise residue contacts. Thus, we show from these experiments that the protein contact map is insufficient for sector extraction and that protein sectors extracted from sequence coevolution form densely connected components.

### 3.4 Protein families with high structural similarity do not share similar protein sectors

In the previous sections, we investigated the signals in the sequence coevolution matrix that the spectral clustering model was extracting to define protein sectors. Now, we turn to the exploration of what protein sectors may reveal about the evolutionary history of a protein family. In Section 3.2, we observed that protein sectors have clear tertiary structural properties even though sector identification is performed independently of the three-dimensional protein structure information. In this section, we further explore whether protein sectors are correlated between protein families with high structural similarity.

Structurally-related protein families do not share sequence identity, yet their three-dimensional protein structures are strikingly similar. For example, the 3AK8 and 3BT5 protein families (aligned in Figure 4DC-DD) share a similar spatial pattern of alpha helices from their N to C-terminus. Likewise, the sequence coevolution matrices of the aligned pair (Figures 4DA-DB) also share similarities due to the relatedness between their three-dimensional structural contacts.

**Figure 4:**
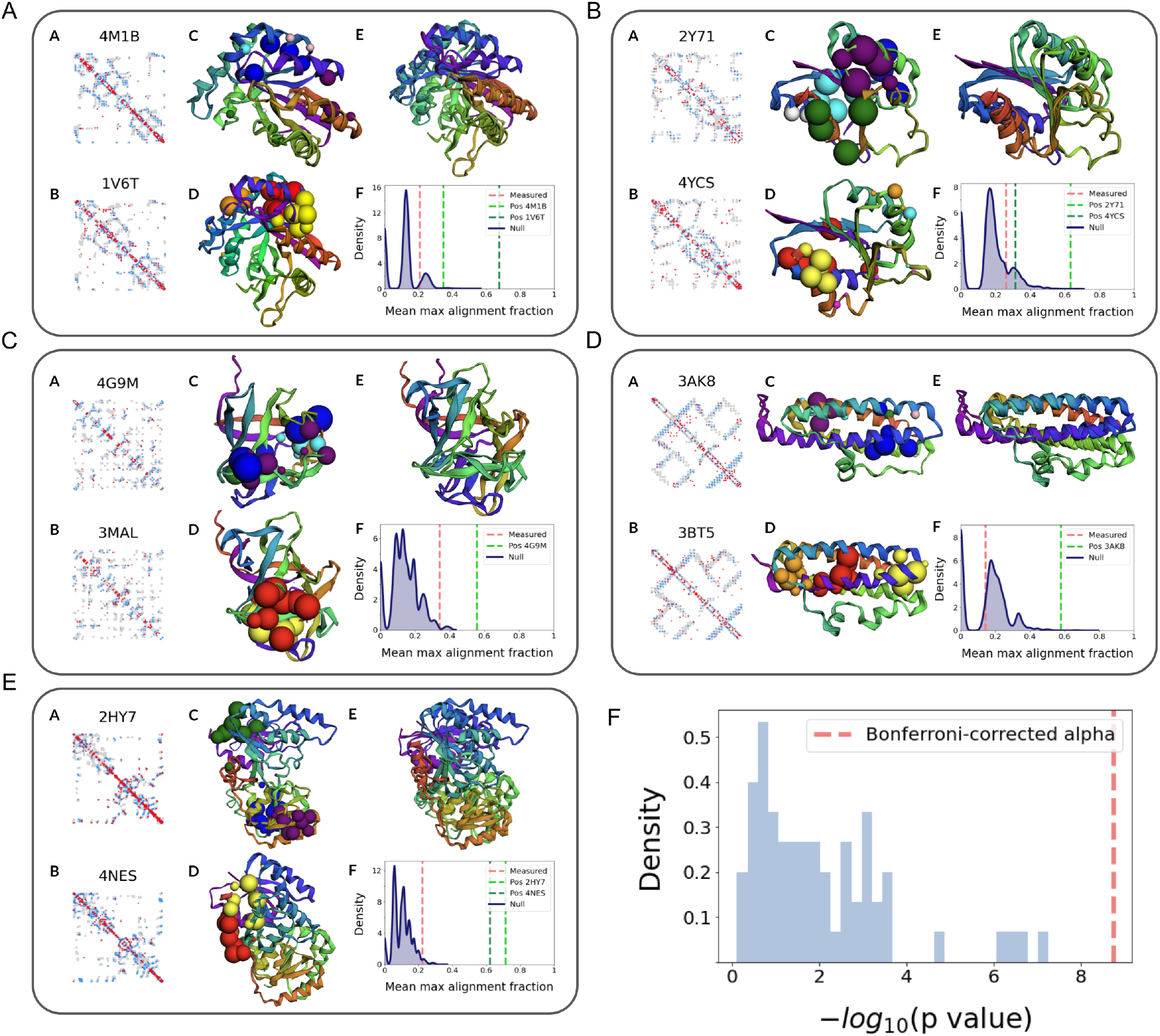
Protein sectors of structurally-related protein families are not equivalent. (A-F) Comparison of protein sectors (AC-AD, BC-BD, CC-CD, DC-DD, EC-ED) between 5 pairs of protein families with high structural similarity (PDB alignments shown in AE-EE). The MSA for a given protein family is trimmed to contain only residues that are structurally-aligned with its paired protein family. Trimmed MSA’s are used to generate sequence coevolution matrices (AA-EA, AB-EB), which is overlaid on its corresponding protein contact map (grey dots). Blue dots denote true positive signals and red dots denote false positive signals. Mean maximum alignment fractions (AF-EF) for extracted sectors (measured; red), positive control sectors (green), and 100 randomly-generated sectors (null; blue). Positive controls are generated for protein families with*≥* 2000 aligned sequences by comparing extracted sectors from mutually exclusive bootstrapped halves of the MSA. (F) Sector alignment significance across 63 pairs of structurally-related protein families. The p value is computed by comparing the measured mean maximum alignment fraction to its null distribution. We use a Bonferroni-corrected alpha value, where *Α* = 0.01.

We examine the similarity between protein sectors across 63 pairs of structurally-related protein families. We have trimmed the MSA’s between pairs of proteins, so we are only considering the aligned residues. Visually, we observe that sectors from 5 of these pairs do not overlap (Figures 4A-E). We can quantify this overlap across 63 pairs of proteins by computing the mean maximum alignment fraction (described in “Methods and Materials”). For example, we compute the mean maximum alignment fraction between the 3AK8 and 3BT5 protein families (Figure 4DF; red dotted line). Compared to a null distribution of randomly-generated sectors, the measured metric is not significantly greater than expected by chance. Moreover, we can also compare the measured metric to a positive control (green dotted line) for 3AK8, in which we compare extracted sectors from mutually exclusive boot-strapped halves of the MSA. The mean maximum alignment fraction for the positive control is nearby 0.6, which is over thrice the size of the measured metric. Thus, we can conclude there is no significant sector overlap between this example pair of proteins.

Continuing this analysis across 63 pairs of structurally-related protein families, we can obtain a distribution of p values (Figure 4F) that describe the deviation of the measured metric between pairs from their null distributions. None of the protein sectors from structurally-related protein families have significant overlap (p value*≤ Α*_*Bonf*_, where *Α* = 0.01), which reveals that pairs of structurally-related protein families do not share similar protein sectors. This suggests that we can use protein sectors to differentiate between pairs of structurally-related protein families.

## 4 Discussion

The massive increase in sequencing data has inspired efforts to extract the biological features of a protein that may be under evolutionary pressure from the sequence data alone. Analyses on single-residue conservation and pairwise sequence coevolution have shown considerable promise in illuminating physical interactions and functional relationships between residues in protein families. These findings have motivated further analysis on higher-order correlations between sequence positions in an effort to further understand how the MSA encodes the conserved biological properties of a protein family. Here, we carry out this analysis using sequence coevolution data predicted by a global statistical model for 120 protein families.

We propose a robust and generalizable spectral clustering model to extract networks of coevolving residues – termed “protein sectors” – for each protein family. We found that protein sectors are structurally connected in three-dimensional space, despite the fact that spectral clustering analysis on the MSA is independent of the three-dimensional protein crystal structure. We show that protein sectors are extracted from a subset of densely connected components in the sequence coevolution matrix, a large number of which are absent from the pairwise residue contact graph that is constructed from the protein crystal structure. Thus, we hypothesize that the sequence coevolution matrix may reveal residue networks consisting of paired residues that are not directly connected in structure but are nonetheless evolutionarily coupled. This reveals a reserve of higher-order evolutionary information previously unnoticed in the sequence coevolution matrix.

The finding that groups of coevolving amino acids may be extracted from the sequence coevolution matrix leads to the question: What part of the evolutionary histories for a given protein family do these protein sectors indicate? We begin this exploration by comparing the protein sectors extracted from structurally-related protein families. Interestingly, we discovered that the extracted sectors of structurally-related protein families do not coincide. Although individual sectors form structurally connected entities, they do not appear to be direct features of the protein structure, suggesting that the protein sectors can help us to differentiate between structurally-related protein families. These findings further support our earlier conclusion that the sequence coevolution matrix contains untapped evolutionary information, which can be valued separately from structure prediction.

We have yet to explore the phylogenetic and functional implications of protein sectors. Namely, we would like to explore whether protein sectors correlate to functional domains in a protein, or whether they reveal groups of residues that diverged in the phylogenetic history of a protein family. Regardless of what protein sectors may represent, the finding that protein sectors constitute a set of densely connected components in the sequence coevolution matrix that are otherwise unattainable from the pairwise contacts graph suggests that there exists networks – not simply pairs – of coevolving amino acids. This is a finding that challenges our basic understanding on the design of natural proteins and urges further investigation on deciphering the largely unknown evolutionary histories of protein families.

## 5 Methods and Materials

### 5.1 Multiple Sequence Alignment

A nonredundant set of 9846 protein chains was collected by the PISCES server [16], accessed on August 2, 2016 [17]. For every protein chain, an MSA was constructed by the HHblits program [18] that was run against the UniProt database [19]. Protein sequences were drawn from hundreds to thousands of organisms, the majority of which are prokaryotes such as *Escherichia coli*. However, eurkaryotic organisms, such as human, mouse, rat, cow, dog, chicken, and zebrafish, are also included. The generated MSA was filtered by HHfilter to exclude similar sequences at 90% identity cutoff as well as sequences with coverage *<* 75%. Positions in the MSA that contain *>* 25% gaps across the sequences were also eliminated. Lastly, in these experiments, only multiple sequence alignments composed of at least 1000 sequences are considered in order to yield more accurate coevolution predictions via GREM-LIN.From HHsearch, we acquired E-values for pairs of the remaining protein families. We removed pairs with an E-values*≥* 1. We also replaced any redundant protein pairs (in which groups of proteins are inter-related with E-values*≥* 1) with a randomly-selected pair to represent the group. This leaves us with 63 pairs, with 120 protein genes in total. We used TM-align to align sequences across the 63 pairs of structurally-related protein families.

### 5.2 Protein Sector Extraction

We develop a spectral clustering algorithm that can predict protein sectors from the MSA of a given protein family. An overview of the procedure (Figure 5) involves computing a similarity matrix to describe the pairwise distances between positions in the protein sequence, transforming the similarity matrix from a high-dimensional to a low-dimensional space, and clustering the reduced data. These clusters will ultimately define our protein sectors.

**Figure 5:**
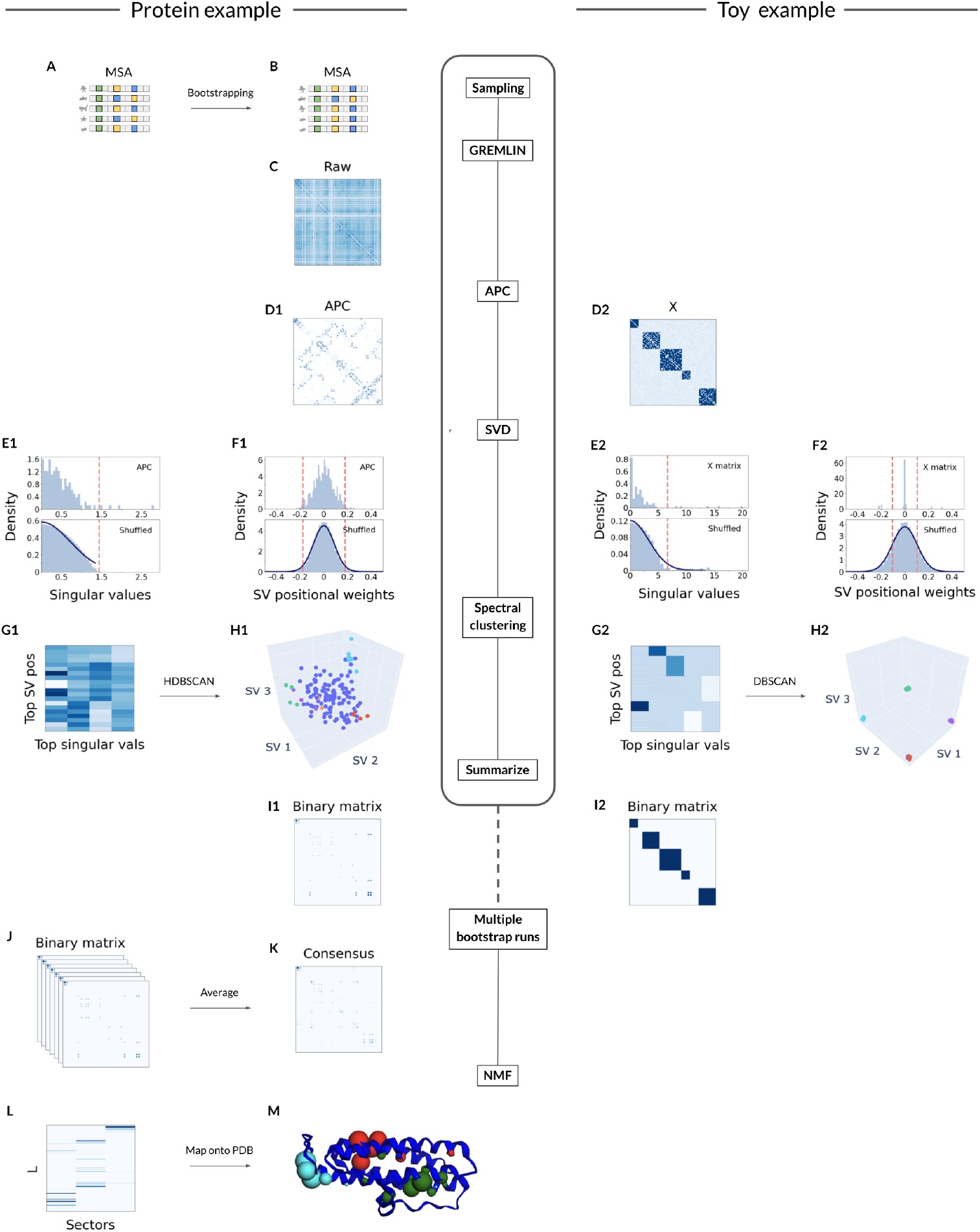
Sector extraction models for protein and toy examples, in parallel. Sequences in the MSA (A) of the 3AK8 protein family are sampled with replacement. GREMLIN models the bootstrapped MSA (B) to estimate the sequence coevolution matrix (C). The AP-corrected coevolution matrix (D1) can be likened to a toy adjacency matrix *X* (D2). We apply SVD to the coevolution (D1) and adjacency matrices (D2) to extract singular values (E1, E2) and vectors (F1, F2), which we compare to a null distribution. We extract the top singular values and the indices corresponding to the top singular vector (SV) positional weights, resulting in a low-dimensional projection of the decomposed coevolution (G1) and adjacency matrices (G2). We perform HDBSCAN and DBSCAN on the projected matrices (G1, G2), respectively, and the resultant clusters are plotted (H1, H2) (top SV positions are in colored dots other than blue) and summarized in binary matrices (I1, I2). The procedure from B-C and D1-I1 is repeated for multiple bootstrap runs. Binary matrices across runs are averaged (J) to yield a consensus matrix (K), which is decomposed by NMF clustering (L). Protein sectors are mapped onto the 3AK8 PDB structure (M) as differently-colored spheres. Sphere size at a given position corresponds to the NMF probability that the residue belongs to a specific sector.

To compute the similarity matrix, GREMLIN models the MSA using a Markov Random Field and implements a pseudo-likelihood approach to optimize its learned parameters. The AP-correction of the weights *w* parameter (Figure 5C) yields the sequence coevolution matrix (Figure 5, D1), which will serve as the similarity matrix. We decompose the coevolution matrix into its singular values and left-singular vectors (Figures 5, E1-F1), which separates functionally significant coevolution signals from those that could be the result of statistical noise. Many singular values and left-singular vector elements of low magnitude can be attributed to statistical noise, so we only interpret the top singular values and the indices that correspond to the top left-singular vector elements. This process leaves us with a projection of the similarity matrix onto lower dimensions (Figure 5, G1). Lastly, we perform hierarchical clustering (HDBSCAN) on the projected low-dimensional coevolution matrix to extract clusters, which correspond directly to the protein sectors (Figure 5, H1).

The spectral clustering model extracts protein sectors given a set of aligned sequences for a protein family. However, the model is sensitive to slight deviations in the sequence coevolution matrix, which can easily arise from sampling variations. Thus, we employ bootstrapping to increase the robustness of the spectral clustering model. We sample sequences with replacement from the MSA to simulate noise from limited sampling of sequences (Figures 5A-B). We extract the protein sectors from each of the sampled multiple sequence alignments by using the spectral clustering model (Figures 5B - H1), and we summarize these residue groupings in a binary matrix (Figures 5, I1). Averaging the binary matrices from each bootstrap run (Figure 1J), we compute a consensus matrix that summarizes the results across all iterations through the spectral clustering model (Figure 5K).

In the final step, we leverage non-negative matrix factorization (NMF) to obtain a consensus clustering across the varied clustering results generated from the bootstrap runs. We use NMF clustering with a softmax activation function to decompose the consensus matrix into two non-negative matrices *W* and *H* (matrix *H* is shown in Figure 5L). The *i, j*-th element of *H* can be interpreted as the probability that residue *i* is part of sector *j*. These NMF probabilities of the decomposed consensus matrix are the residue groupings that define the protein sectors (Figure 5M).

In parallel, we also perform the spectral clustering model on a toy example in Figure 5, D2-I2. The adjacency matrix of the toy example (Figure 5, D2) clearly denotes 5 clusters, which are reconstructed in the resultant binary matrix (Figure 5, I2) that is returned by the model. Based on these results, we expect that the spectral clustering model can reliably extract protein sectors in the multiple sequence alignment.

### 5.3 Structural Connectivity of Protein Sectors

We explore the relationship between a residue’s probability of sector membership and its physical relativity to the sector’s center. To measure this relationship, we first obtain a centroid for each sector.

#### 5.3.1 Sector Centroid

The centroid (*x, y, z*)_*j*_for sector *j* is computed by taking the weighted average of the coordinates of the residues involved in the sector:

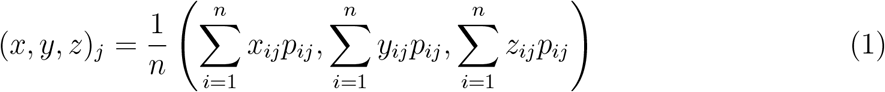

Where *n* is the number of residues that are part of sector *j*, (*x*_*i*_, *y*_*i*_, *z*_*i*_) are the three-dimensional coordinates for residue *i* as given in the PDB structure, and *p*_*i*_is the probability that residue *i* is part of sector *j*. The probability *p*_*i*_is taken from the *j, i*-th element of the *H* matrix obtained in non-negative matrix factorization (NMF).

#### 5.3.2 Distance from Sector Centroid

Having computed the centroid for sector *j*, we can calculate the distance in Angstroms of each residue *i* from the centroid, where residue *i* has a probability *p*_*i*_*≥* 0.01 of belonging to sector *j*.

### 5.4 Densely Connected Components

We investigate whether residue contact information alone is sufficient to produce the same protein sectors as extracted as from the GREMLIN-generated coevolution matrix.

#### 5.4.1 Contacts-Generated Sectors

We compare the graph connectivity of protein sectors generated using the binary contact map as the similarity matrix with the graph connectivity of protein sectors generated using the sequence coevolution matrix.

To obtain the binary contact map, we use the tool ConFind [26]. ConFind computes *cf* scores for each position pair in a protein family. Position pairs with *cf ≥* 0.01 are considered to be in contact and are expected to co-vary evolutionarily. In computing the *L ×L* binary contact map, an element *i, j* was set to 1 if residue pairs *i* and *j* had a ConFind score greater than 0.01.

#### 5.4.2 Scaled Beta Index

The beta index *β* measures the level of connectivity in a graph network:

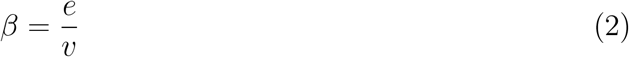

where *e* is the number of edges and *v* is the number of nodes, i.e. the number of residues in a sector. Simple networks have a beta index less than one, connected networks with one cycle have a beta index equal to one, and complex networks have a beta index greater than one.

Given a protein sector, which we can interpret as a graph network, we use the scaled beta index *β*′ to measure the level of connectivity of the largest connected component in the network while taking into account the fraction of residues (i.e. nodes) within the sector that is participating in the largest connected component. We compute the scaled beta index *β*′ by taking the product of the following two elements:

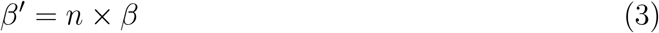

where *n* is the fraction of nodes participating in the largest connected component in the graph, and *β* is the level of connectivity for the largest connected component in the graph network.

#### 5.4.3 Extracting Top Coevolution Signals

To ensure that we were making fair network comparisons between contacts- and coevolution-generated sectors, we extracted the same number of coevolution signals as we did contacts. The contact map is a binary classification of 1’s and 0’s, so the number of extracted contacts is a fixed number *n*. We defined a threshold such that we extracted the *n*-largest elements from the sequence coevolution matrix.

### 5.5 Protein Sectors and Structurally-Related Protein Families

We would like to compare the extracted protein sectors of pairs of structurally-related but discrete protein families. We introduce a metric designed to measure the overlap between two sets of protein sectors: mean maximum alignment fraction *M*.

#### 5.5.1 Mean Maximum Alignment Fraction

We compute the mean maximum alignment fraction *M* to measure the overlap between the protein sectors in protein *A* and the protein sectors in protein *B*:

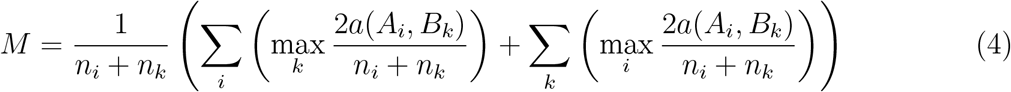

where *A*_*i*_is a sector *i* in protein *A, B*_*k*_is a sector *k* in protein *B, n*_*i*_is the number of residues in sector *i*, and *n*_*k*_is the number of residues in sector *k*. The operator *a*(*A*_*i*_, *B*_*k*_) is the number of aligned residues between *A*_*i*_and *B*_*k*_.

The mean maximum alignment fraction can range from 0 to 1, where 0 describes sectors with no overlapping residues and 1 describes identical sectors.

## 6 Supplementary Methods

### 6.1 Spectral Clustering Model

Spectral clustering allows us to generate protein sectors from the multiple sequence alignments of a given protein family (Figure 5). The process involves 3 steps: computing an *L× L* (*L* is the protein length) similarity matrix, projecting the matrix onto a low-dimensional space, and creating clusters, which will ultimately define our protein sectors.

Sections 6.1.1 and 6.1.2 describe how we compute the similarity matrix. We use the GREMLIN Learning Algorithm to model the multiple sequence alignment (MSA) for each protein family and extract its learned parameters. The average product correction (APC) [20] of the Frobenius norm of the weights *w* parameter represents the sequence coevolution matrix for a given MSA, which will serve as the similarity matrix.

Section 6.1.3 describes how we separate the functionally significant coevolution signals from those that could be the result of statistical noise. We employ singular value decomposition (SVD) to extract the singular values and left-singular vectors of the sequence coevolution matrix. Many singular values and left-singular vector elements of low magnitude can be attributed to statistical noise, so we only interpret the top singular values and the indices that correspond to the top left-singular vector elements. This process leaves us with a projection of a condensed similarity matrix onto lower dimensions.

Lastly, section 6.1.4 describes how we cluster a condensed version of the low-dimensional projected data. We use Hierarchical Density-based Spatial Clustering of Applications with Noise (HDBSCAN) [21], which is a deterministic hierarchical clustering algorithm. The returned clusters correspond directly to the “protein sectors”.

#### 6.1.1 The GREMLIN Learning Algorithm

Lapedes et al. (1998, 1999) proposed a Markov Random Field (MRF) that accounts for single-position conservation and pairwise co-evolution. The MRF has the following form, in which the probability of a given sequence *X* is:

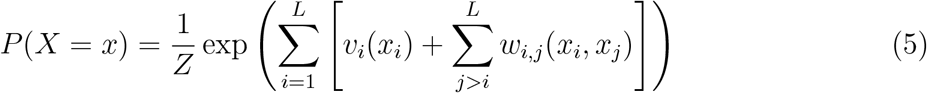

A model is learned for each multiple sequence alignment of a protein family. The random variables *x*_*i*_is the amino acid residue at position *i, v*_*i*_is a set of parameters that encodes the individual propensity for each amino acid at position *i* of the protein, and *w*_*i,j*_is a set of parameters that model the statistical coupling in amino acid propensities between positions *i* and *j. Z* is the partition function, normalizing the probabilities such that they sum to 1 [11, 22].

The parameters *v* and *w* are optimized to maximize the probability of the given sequence. However, solving Equation 5 requires us to consider all possible sequences of length *L* (re-sulting in 20*L* sequences total) in order to compute the partition function *Z*. This problem is computationally intractable for computing a single likelihood, let alone optimizing for *v* and *w*.

To resolve this, the GREMLIN (Generative REgularized ModeLs of proteINs) Learning Algorithm was proposed. GREMLIN implements a pseudo-likelihood approach to optimize *v* and *w* [10, 23].

Given a set of aligned protein sequences (i.e. the multiple sequence alignment, MSA), GREMLIN learns the parameters by optimizing the pseudo-likelihood of θ (*v* and *w*):

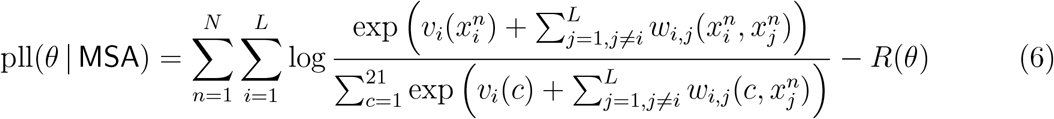

It has been shown that the *θ* that maximizes the pseudo-likelihood pll(*θ* | MSA) (Equation 6) has the equivalent notion to the *θ* that maximizes the likelihood *L*(*θ* | MSA) (Equation 5) under certain conditions and assumptions, see [24]. *N* is the number of sequences in the multiple sequence alignment, *L* is the sequence length, 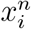is the amino acid residue at position *i* in sequence *n*, and *c* is the 21 characters (20 amino acid residues and 1 gap). *R* is the regularization term that promotes a sparse network:

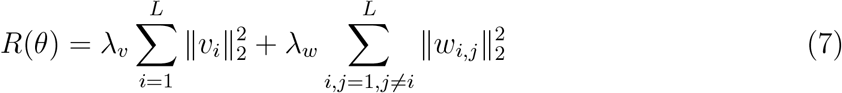

where *λ*_*v*_is 0.01 and *λ*_*w*_is 0.2(*L−1*). The pseudo-likelihood models the conditional distributions of the original joint distribution instead of the joint distribution itself. The global partition function *Z* in Equation 5 has been replaced with a local *Z* in Equation 6. The local partition function is concave in *v* and *w*, so the learning procedure is consistent and easy to maximize.

#### 6.1.2 Obtaining Sequence Coevolution

We use GREMLIN to obtain the *L × L* sequence coevolution matrix for each multiple sequence alignment (MSA) in our protein dataset. The coevolution matrix will serve as the similarity matrix in the downstream spectral clustering model.

Computationally, GREMLIN is an autoencoder with a single dense layer between the input and output. The model seeks to minimize the difference between the input (the given MSA) and output (the predicted MSA) by optimizing the weights *w* and bias *v* in the dense layer. We extract the weights *w* from the pseudo-likelihood framework, and we apply an average product correction (APC) to the Frobenius norm of the weights *w*. The resultant *L × L* matrix *w*′ represents the sequence coevolution, in which each entry 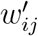is the strength of coevolution between residues *i* and *j*.

Average product correction (APC) is used to disentangle signals caused by structural and functional constraints from background signals caused by random noise and phylogeny [20]. APC is effectively removing the largest eigenvector from the sequence coevolution matrix. This process is necessary to induce a sparser network because the regularization term in Equation 7 is not a constraint.

#### 6.1.3 Spectral Cleaning

Inspection of the sequence coevolution matrix indicates that strong coevolution signals are not simply dominated by proximity in primary structure. Many positions show weak coevolution to neighboring positions but significant coevolution to positions that are more distant along the sequence.

However, before we can extract the residue protein sectors from the sequence coevolution matrix, we have to first separate the functionally significant coevolution signals of *w*′ from correlations that could have arisen from statistical noise due to limited sampling of sequences. Singular value decomposition (SVD) provides a method to partially sort out the different contributions to the coevolution signals. In SVD, the sequence coevolution matrix *w*′ is decomposed into three matrices: *w*′ = *U* σ*V* ∗. The diagonal entries *σ*_*i*_of σ are the singular values of *w*′, the columns of *U* are the left-singular vectors of *w*′, and the columns of *V* are the right-singular vectors of *w*′.

The singular values *σ*_*i*_and left-singular vectors *U* are related to the eigenvalues and eigenvectors of *w*′, respectively. In fact, the singular values are equal to the absolute value of the eigenvalues because the sequence coevolution matrix is symmetric. We choose to decompose the sequence coevolution matrix using SVD rather than eigenvalue decomposition because we would like to sort the eigenvectors by their absolute eigenvalues for easier interpretability. The spectrum of *w*′ is composed of *L* singular values, the lowest of which can be attributed to statistical noise since shuffled sequence coevolution matrices preserving local residue interactions show singular values of similar magnitude. We extract the top singular values by comparing them to generated singular values from 200 random simulations. The randomization procedure involves computing the singular values of sequence coevolution matrices shuffled along the off-diagonal axes. This method of shuffling retains the local residues that are in contact with each other by virtue of protein folding. The randomized singular value distribution is fit by a half-normal distribution model. The half-normal distribution has mean *µ* = 0 and standard deviation 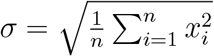where *x*_*i*_is the *i*-th singular value. We extract the singular values of *w*′ that are greater than two standard deviations above the mean.

Thus, we only interpret the top singular values and their corresponding left-singular vectors. Furthermore, we continue to extract only the top indices of the left-singular vectors. We again compare the left-singular vector weights of *w*′ to those generated from 200 random simulations. The sequence coevolution matrices are similarly shuffled along the off-diagonal axes. The left-singular vector weights distribution is fitted by a Gaussian distribution. Two standard deviations above and below the mean are used as significance thresholds for determining the top index positions. We extract the left-singular vector elements that correspond to these top index positions.

This process of spectral cleaning leaves us with a low-dimensional projection of the original sequence coevolution matrix *w*′. The low-dimensional projected data 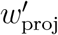has dimensions *P × S*, where *S* is the number of top singular values and *P* is the number of top index positions.

#### 6.1.4 Generate Clusters

We cluster the low-dimensional projected data using Hierarchical Density-based Spatial Clustering of Applications with Noise (HDBSCAN) [21]. This hierarchical clustering algorithm transforms the space according to density, builds a minimum spanning tree of the distance weighted graph, constructs a cluster hierarchy of connected components, condense the cluster hierarchy based on minimum cluster size, and extracts the stable clusters from the condensed tree.

In contrast to typical clustering algorithms, HDBSCAN is more robust to noise because it takes into account the density of the data. Moreover, we found that clustering 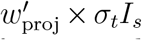 yields more informative clustering results, so we apply HDBSCAN to this transformed and condensed low-dimensional projected data. Here, *σ*_*t*_*I*_*s*_refers to a diagonal matrix with entries that correspond to the top singular values, and 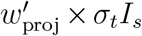is a (*P S*) dimensional matrix in which each column of 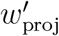has been multiplied by its corresponding singular value in *σ*_*t*_*I*_*s*_. The extracted clusters from HDBSCAN represent our protein sectors.

### 6.2 Model Robustness

The spectral clustering model allows us to extract “protein sectors” given a set of aligned sequences for a protein family. However, the model is sensitive to slight deviations in the sequence coevolution matrix, which can easily arise from sampling variations. The cause of the model’s sensitivity stems from the spectral cleaning of left-singular vector index positions. The inclusion or exclusion of certain positions with vector elements near the boundaries of thresholds can heavily impact downstream clustering.

#### 6.2.1 Bootstrapping

We perform bootstrapping to increase the robustness of the spectral clustering model. We sample sequences with replacement from the multiple sequence alignment (MSA) to simulate noise from limited sampling of sequences. We extract the protein sectors from each of the sub-sampled MSA’s by using the spectral clustering model.

To increase the speed of GREMLIN, we initialize the model parameters *w* and *v* for each sub-sampled MSA with the learned weights and biases from a GREMLIN model for the full MSA that was optimized over 1000 iterations. Then, we only need to optimize the GREMLIN model for sub-sampled MSA’s for 100 iterations.

#### 6.2.2 Softmax Non-Negative Matrix Factorization

Non-negative matrix factorization (NMF) factorizes a matrix *V* into two non-negative matrices *W* and *H*. The objective function is:

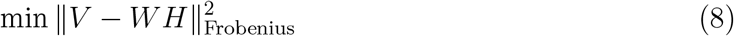

with the constraints *H ≥* 0, *W≥* 0. The Forbenius norm is taken as the square root of the sum of the squared elements. NMF has an inherent clustering property in that matrix *H* details cluster membership: if *H*_*kj*_*> H*_*ij*_for all *I k*, then the data at the *j*th column of *V* belongs to the *k*th cluster.

We can leverage NMF clustering to obtain a consensus clustering across the varied clustering results generated from bootstrap runs. To do this, we first obtain an *L × L* binary “common” matrix *C* for each bootstrap run in which each element *C*_*i,j*_describes whether residues *i* and *j* are in the same sector as determined by HDBSCAN. After *b* bootstrap runs, we can obtain a consensus matrix, which is the average over these common matrices: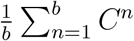

We perform NMF clustering on the consensus matrix to obtain two non-negative matrices *W* and *H*. We implement the algorithm in TensorFlow, using the objective function given in Equation 8. The objective function is minimized with an alternating minimization of *W* and *H*. Additionally, we apply a softmax activation function along the rows of *W* and the columns of *H* to normalize the output to a probability distribution over predicted output classes. Thus, the *i, j*-th element of *H* can be interpreted as the probability that residue *j* is part of sector *i*.

#### 6.2.3 Selecting the Number of Clusters

Non-negative matrix factorization requires you to select the number of components *k* (i.e. the number of protein sectors) that you would like to decompose the matrix into. We use the elbow method to select the optimal number of components *k*. The idea of the elbow method is to first perform NMF clustering on the consensus matrix 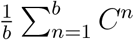 for a range of values of *k*, let’s say from *k* = 1 to *k* = 10. For each value of *k*, we calculate the sum of squared errors (SSE), which is computed as:

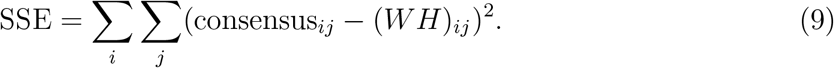

We create a plot to visualize the relationship between the SSE and the number of components *k*. If the plot looks like an arm, then the “elbow” of the arm is the optimal value of *k*. Intuitively, increasing *k* will naturally improve the fit and have a lower SSE, but at some point this will result in over-fitting, and the “elbow” reflects this inflection point. Using this heuristic in mathematical optimization allows us to choose a point where diminishing returns are no longer worth the additional cost.

Selecting the “elbow” of the plot is generally a subjective task. We can automate this by selecting the number of components *k* that corresponds to the maximum elbow strength *S*. To calculate elbow strength *S*, we first compute the Δ scores for each *k*, which is equal to the first-order differences between the sum of square errors: *SSE*_*k*1_*SSE*_*k*−1_. Then, we compute the Δ′ scores for each *k*, which is equal to the second-order differences between the sum of square errors: Δ_*k*1_Δ_*k*_. Lastly, the elbow strength *S* for each *k* is computed as the difference between Δ′ and Δ scores at 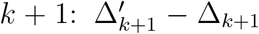. The optimum number of components *k* is the value of *k* that maximizes the strength *S*.

## 7 Code Availability

Code can be found at the GitHub repository: https://github.com/cs975/protein_sectors. We have included an interactive notebook where readers can investigate the protein sectors for any protein family.

